# Recognition and localization of maize leaves in RGB images based on Point-Line Net

**DOI:** 10.1101/2024.01.08.574560

**Authors:** Bingwen Liu, Jianye Chang, Dengfeng Hou, Dengao Li, Jue Ruan

**Affiliations:** College of Computer Science and Technology (College of Data Science), Taiyuan University of Technology, Taiyuan, 030024, China; Shenzhen Branch, Guangdong Laboratory of Lingnan Modern Agriculture, Genome Analysis Laboratory of the Ministry of Agriculture and Rural Affairs, Agricultural Genomics Institute at Shenzhen, Chinese Academy of Agricultural Sciences, Shenzhen 518120, China

**Keywords:** Maize, Deep learning, Object detection, keypoint detection

## Abstract

Plant phenotype detection plays a crucial role in understanding and studying plant biology, agriculture, and ecology. It involves the quantification and analysis of various physical traits and characteristics of plants, such as plant height, leaf shape, angle, number, and growth trajectory. By accurately detecting and measuring these phenotypic traits, researchers can gain insights into plant growth, development, stress tolerance, and the influence of environmental factors. Among these phenotypic information, the number of leaves and growth trajectory of the plant are more accessible. Nonetheless, obtaining these information is labor-intensive and financially demanding. With the rapid development of computer vision technology and artificial intelligence, using maize field images to fully analyze plant-related information such as growth trajectory and number of leaves can greatly eliminate repetitive labor work and enhance the efficiency of plant breeding. However, the application of deep learning methods still faces challenges due to the serious occlusion problem and complex background of field plant images. In this study, we developed a deep learning method called Point-Line Net, which is based on the Mask R-CNN framework, to automatically recognize maize field images and determine the number and growth trajectory of leaves and roots. The experimental results demonstrate that the object detection accuracy (mAP) of our Point-Line Net can reach 81.5%. Moreover, to describe the position and growth of leaves and roots, we introduced a new lightweight “keypoint” detection branch that achieved 33.5 using our custom distance verification index. Overall, these findings provide valuable insights for future field plant phenotype detection, particularly for the datasets with dot and line annotations.

## Introduction

Maize, as an essential cash and food crop worldwide, not only provides a significant amount of food and feed for humans but also finds extensive applications in industries, bio-energy and other sectors. The study of maize phenotypes encompasses the examination of external morphological and physiological characteristics, including plant height, leaf number, leaf morphology, and more. By investigating maize phenotypes, vital data and information for molecular breeding can be obtained, effectively advancing the modernization and efficiency of maize breeding. Moreover, this research plays a crucial role in promoting the development of maize industry and improving maize production efficiency and quality. Nevertheless, acquiring most phenotypic characteristics requires substantial human, material, and financial resources. This limitation significantly impacts the research process regarding plant phenotypic characteristics. For instance, obtaining the precise number of maize leaves often demands repetitive statistical corrections to mitigate errors, leading to a significant waste of resources. Consequently, automated and reliable techniques are indispensable for the detection of maize field phenotypes to enhance efficiency and reduce resource consumption in maize phenotype research. In recent years, the field of plant phenotype detection has witnessed a notable advancement, thanks to the rapid development of computer vision technology, especially deep learning. Deep learning, a branch of machine learning, utilizes artificial neural networks to model and solve complex problems. By training a neural network with a vast amount of data, the network can learn and enhance its performance over time through a process known as back-propagation. Among various deep learning algorithms, classification algorithms are widely employed to categorize input data into one or more classes using deep neural networks. For instance, (1) proposed a convolutional neural network (CNN) to enhance the classification of plant leaf diseases. (2) introduced a new and lightweight architecture called DLMC-Net, which demonstrated the capability of real-time agricultural applications in detecting plant leaf diseases across multiple crops. (3) developed a deep-learning-based classification method to evaluate the extent of maize lodging using RGB and multi-spectral images captured by unmanned aerial vehicles (UAVs). They analyzed the characteristic variations exhibited in RGB and multi-spectral images, corresponding to three different lodging extents. Semantic segmentation is a computer vision technique that aims to assign a semantic label to each pixel in an image, thereby segmenting the image into meaningful regions based on their semantic content, such as objects, backgrounds, or other important features (4). Leaf segmentation, in particular, has always posed a challenge due to the overlapping nature of leaves and the complex background. To address this, a novel convolutional neural network named LS-Net has been proposed specifically for leaf segmentation in rosette plants (5). The experiment conducted by Deb et al. Utilized a dataset of over 2010 images from the plant phenotyping (CVPPP) and KOMATSUNA datasets. In another study, (6) examined the accuracy differences of the U-Net model when applied to images of various maize varieties, different stages of tasseling, and different spatial resolutions. In addition to the aforementioned techniques, object detection represents another crucial computer vision approach that enables the identification of objects in images or videos by drawing a bounding box around them. Various popular algorithms have been developed for object detection, including Faster R-CNN (7), YOLO (You Only Look Once) (8), and SSD (Single Shot Detector) (9). These algorithms typically combine feature extraction, region proposal, and classification method to achieve accurate object detection. In the field of agriculture, object detection algorithms have emerged as vital tools for the detection and classification of plants. (10) introduced a novel benchmark for YOLO object detectors in the context of multi-class weed detection in cotton production systems. Their proposed benchmark showcases a more accurate and efficient approach compared to traditional manual weed detection methods. (11) developed an automatic detection method based on the improved Mask R-CNN framework. This method improved efficiency and reduced the high cost associated with seedlings sorting in the raising process of hydroponic lettuce seedlings. Additionally, some researchers have leveraged an improved Cascade R-CNN architecture known as RiceRes2Net (12) to detect rice panicles and identify growth stages in complex field environments (13). This approach contributes to advancing the understanding and management of rice cultivation by accurately identifying and tracking panicles development. Although deep learning methods have been widely applied to plant phenotype detection, none of them have been utilized for field maize leaf detection and precise localization of leaf veins. In this paper, we will present our approach to address this issue.

## Methods

### Maize dataset

The maize dataset used in this study, as shown in Figure 1, was provided by the Agricultural Genomics Institute, Chinese Academy of Agriculture Sciences. Our team has constructed a comprehensive maize image phenotype dataset, which was collected using handheld cameras and unmanned aerial vehicles(UAV) at multi angles and time intervals. The dataset encompasses more than 32,000 high-resolution RGB images, of which over 18,000 leaves and roots were manually labeled using Labelme software and all maize images were divided into three distinct periods based on growth cycles. For this study, due to noteworthy disparities in morphology and background between maize images captured by cameras and those obtained via UAV, we decided to exclusively analyze and process the maize images acquired through cameras to attain superior results.Nevertheless, there exists immense potential to extract additional insights from the multiple-angle and multi-time series data to make further progress, which we will discuss this in Section Conclusions. Compared to currently widely used plant phenotype datasets, which are primarily annotated with segmentation masks or target detection rectangles((14), (15), (16)), our maize dataset takes a novel approach by utilizing dotted-line annotation to depict leaf and root veins, as illustrated in Figure 1. The method offers the advantage of conveying a more precise representation of these veins. when target positions and contours are labeled using segmentation masks, semantic segmentation or instance segmentation algorithms (17) often generate extraneous pixels. However, not all pixels are required to describe leaf and root veins. In addition, it is inappropriate to use solely rectangular bounding boxes to annotate target positions, as unlike the laboratory environment (14), there are significant occlusion issues with maize leaves in field conditions, especially in later stages. This poses an important challenge that we need to address in our work. Hence, the utilization of dotted line annotation is more suitable for our field-based maize phenotype detection, at the same time, providing researchers with greater challenge.

**Fig. 1.**
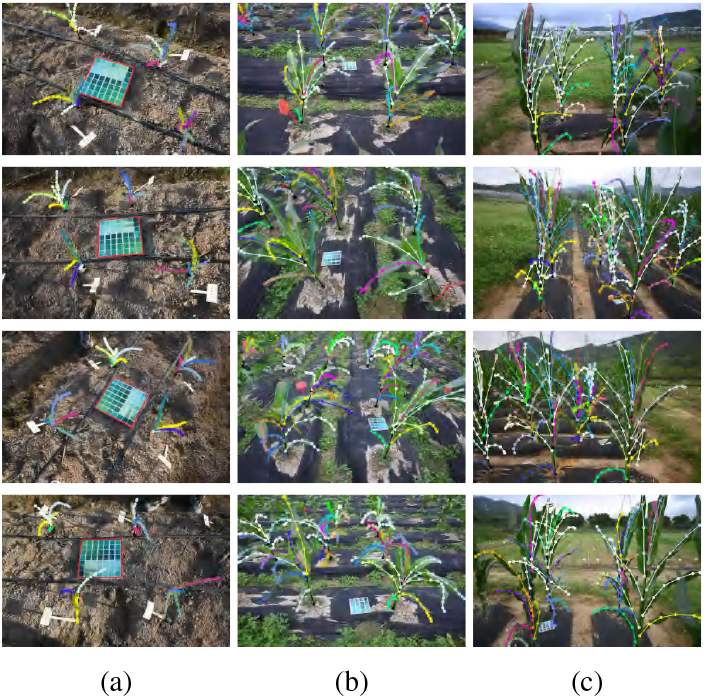
Images at 4 angles of maize dataset with annotations for three growth periods. (a) in the early stage, (b) in the middle stage, (c) in the late stage.

### Dataset processing

The original field images in our dataset have a size of 6016*4016(or 4016*6016). Considering the challenges associated with memory management on available GPUs, we manually downsampled the original images by a factor of 4 to 1504*1004(or 1004*1504) for training and validating. We also downscaled the coordinate in-formation stored in the corresponding annotation file accordingly. This downsampling approach was adopted to address the limitations of memory size while aiming to retain the RGB information as much as possible. The annotation file is formatted as a JSON file containing comprehensive annotation information, including the target category (such as leaf or root), the serial number, and the coordinate information of the annotated points, etc. As mentioned previously, our original dataset does not provide rectangular bounding boxes for target detection considering the desire to solve the target occlusion problem. However, our proposed approach based on Mask R-CNN requires the annotation of actual target detection boxes (18). To address this, we calculate the coordinates of the points marked for each individual target, determining the minimum coordinate values as the upper-left point of the target detection box and the maximum of coordinate values as the lower-right point of the target detection box. We also added a certain offset(20) for each calculated coordinate point to include the full target, as depicted in Figure 2. In this way, the generated rectangular box serves as the ground truth for the training process. Additionally, inspired by the multi-person human keypoint detection task((19), (20)), our proposed approach aims to extend the object detection task to a keypoint detection task and further details will be discussed in Section Modification of keypoint detection branch. To make cluttered raw keypoint annotations more learnable, we uniformly process the dataset using the following method: Firstly, we employ an interpolation algorithm to expand the initially small number of labeled keypoints. Then, for each ground-truth object, we traverse the keypoints contained within it and extract a portion of the them as ground-truth keypoints at equal intervals based on a predetermined ratio(30%).The final data processing results are shown in Figure 2.

**Fig. 2.**
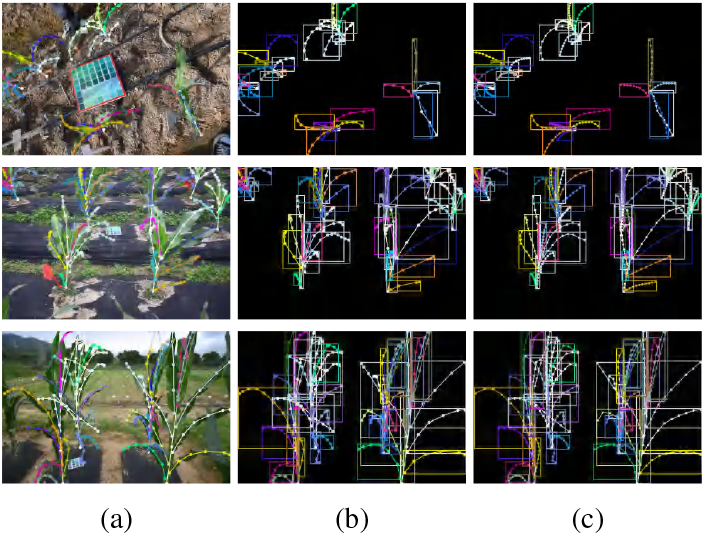
Method of determining ground-truth bounding box. (a) original annotation, transformed bounding box annotation, (c) key-point annotation after interpolation algorithm processing.

### Maize detection

#### Point-Line Net Based Mask R-CNN

Taking into account the distinctive dotted-line annotation in our dataset, we draw a parallel between this annotation and the task of human pose estimation (20). Human pose estimation, often referred to as keypoint detection, is a computer vision task that involves locating and identifying specific points on on a person’s body, such as joints, limbs, and facial features. The objective of keypoint detection is to accurately estimate the 2D or 3D positions of these points in an image or video. Similar to the instance segmentation task, keypoint detection can be categorized into two approaches: top-down and bottom-up, based on the pose formation logic. The top-down approach, which relies on target detection (18), involves first detecting the regions in the image where each instance is presented, and then performing a separate semantic segmentation task or key-point detection on these candidate regions. This approach is constrained by the accuracy of the object detection stage. On the other hand, the bottom-up approach (17) treats instance segmentation or multi-person keypoint detection as a clustering task, grouping pixels or global keypoints into an arbitrary number of target instances within the image. The class of each group is then determined to achieve instance segmentation or keypoint detection. Given the complex background of the field and the challenge of overlapping shading, particularly in later stages, we proposed a improved top-down approach based on Mask R-CNN named Dotted-Line Net, which aims to address these challenges and improve the accuracy of keypoint detection in our dataset. Mask R-CNN is an extension of Faster R-CNN, as depicted in Figure 3. Faster R-CNN consists of two main components: a region proposal network (RPN) and a Fast R-CNN detector. The RPN generates a set of object proposals, which are then inputted into the Fast R-CNN detector for object classification and bounding box regression. The RPN is trained to learn objectness scores and bounding box regressions, while the Fast R-CNN detector is trained to classify the objects within each proposal and refine the bounding boxes. Building upon Faster R-CNN, Mask R-CNN extends the framework by predicting a binary mask or a set of heatmaps for keypoints for each detected object. The mask indicates the pixel-wise location of the object within the image, while the heatmaps indicate the specific locations of keypoints on the object within the image. Unlike Mask R-CNN, which incorporates a third branch for traditional keypoint detection into the network, our approach modifies Faster R-CNN by introducing a branch to predict “keypoints” for each detected object using a single heatmap, as illustrated in Figure 3. Additionally, the proposals generated by the RPN are also utilized in the keypoint head for keypoint detection. Further details of the keypoint detection branch will be presented in Section Modification of keypoint detection branch.

**Fig. 3.**
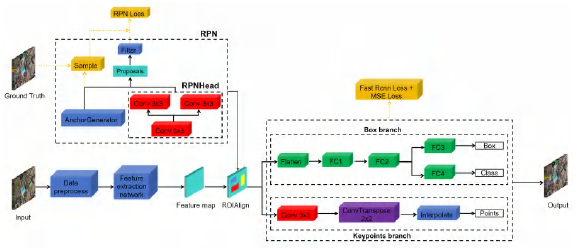
Point-Line Net structure based on Mask R-CNN model.

#### Adjustment of hyper-parameters

We made adjustments to several hyper-parameters and conducted corresponding experiments. Firstly, we increased the number of proposals retained in the RPN before and after Non-Maximum Suppression(NMS) processing, namely RPN_PRE_NMS and RPN_POST_NMS. The numbers were increased from 2000 to 4000 and from 1000 to 2000, respectively. This modification aimed to retain more candidate proposals, taking into consideration that our dataset often features as a larger number of targets, and the distance measured by Intersection over Union (IoU) between targets is closer. The results, as demonstrated in Section Object detection results of maize based on three different original models, indicate that this adjustment yields the best performance for object detection in our model. Additionally, regarding the anchors, we expanded the number of scales to include six different box areas, namely 322, 642, 1282, 2562 and 5122 pixels. This adjustment was experimentally proven to accelerate the convergence of the model and improve its performance, allowing it to adapt to targets of various scales. Furthermore, in the original Faster R-CNN (7), RoIPool is employed to align the proposals obtained from the RPN to the same size. However, this process involved rounding operation, resulting in less precise localization. To overcome this limitation, the authors proposed the RoIAlign method as an alternative to RoIPool, which provides more accurate spatial localization information. We have also incorporated this improvement into our model. Moreover, the size of the pooling operation not only impacts the expressiveness of the network but also affects the accuracy of the RoI (Region of Interest). If the pooling size is too small, such as 7, detailed information within the RoI may be lost, thereby hindering the detection results. Conversely, if the pooling size is too large, such as 28, the feature representation of the RoI becomes too coarse, potentially degrading the detection performance. Taking into consideration the characteristics of out dataset and the aforementioned analysis, we adjusted the output size of RoIAlign to 14, which yielded the optimal detection effect.

#### Soft-NM

NMS(Non-Maximum Suppression) is a commonly used technique in object detection algorithms to remove redundant and overlapping detections, resulting in a final set of bounding boxes with the highest confidence scores. However, NMS has its limitations. It may discard valid detections with lower scores if they overlap with a higher-score detection. Additionally, NMS selects only one detection for each object instance, which can lead to suboptimal performance when multiple detections are valid. To address these limitations, Soft-NMS (21) was introduced, as a modified version of NMS that incorporates a soft weighting scheme. In Soft-NMS, each detection is assigned a weight based on its confidence score and the overlap it has with other detections. The weight of a detection is determined by its score as well as the scores of all other detections that intersect with it. This weighting scheme allows for multiple detections of the same object instance to be retained while suppressing redun-dant detections. We believe that applying Soft-NMS to our maize dataset, which contains a significant amount of occlusion, can yield more accurate and robust results. Our experiments have also confirmed this conjecture.

#### D_IoU

Distance-IoU (22) introduces the concept of considering the distance between the center points of bounding boxes, to better measure the overlap between two closely positioned bounding boxes. This differs from traditional IoU, which only takes into account the ratio of the overlapping area to the union area of two bounding boxes, and is represented as:

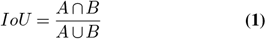

Where *A* and *B* represent the predicted and the ground-truth bounding box respectively. In contrast, Distance-IoU, abbreviated as D_IoU, is more robust to variations in object sizes and aspect ratios. Traditional IoU may exhibit bias towards larger objects, whereas D_IoU can better handle scenarios where objects have different sizes and aspect ratio, as is often observed in our maize dataset. Therefore, D_IoU can provide a better reflection of the similarity and difference between two bounding boxes, avoiding the merging of bounding boxes that are in close proximity or overlapping. To improve the detection accuracy of closely positioned objects, we integrate D_IoU into the NMS processing, which is represented as:

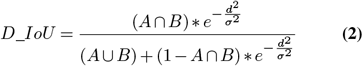

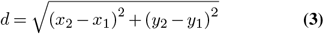

Where (*x*_1_, *y*_1_) and (*x*_2_, *y*_2_) represent the coordinates of the center points of the two bounding boxes respectively, and *σ* denotes an attenuation parameter that controls the degree of influence of distance. In our experiments, *σ* took the value of 0.3.

#### Modification of keypoint detection branch

There are two main approaches to tackle the keypoint detection task: the regression-based approach and the heatmap-based approach. The regression-based approach directly estimates the key-point coordinates (e.g., *x* and *y* coordinates) for each key-point in the image. In this approach, the model is trained to predict the coordinates of each keypoint directly from the input image (23). The output of the model is a a set of coordinates that can be used to visualize the keypoints on the image. While this method is generally straightforward and efficient, it may not achieve the same level of precision as the heatmap-based approach due to the inherent limitations of CNNs in regressing long-distance offsets (24). On the other hand, the heatmap-based approach (25) utilizes a deep neural network to analyze the image or video and generate a heatmap for each keypoint of interest. The heatmap indicates the likelihood of the keypoint being present at each location in the image or video. This approach is known for its higher accuracy but may involve more complex computation and slower processing compared to the regression-based approach. The keypoint detection branch of Mask R-CNN is composed of fully connected layers that receive RoI-Aligned features as input and generate a set of heatmaps, with each heatmap corresponding to a specific keypoint. However, as discussed in Section Maize dataset, the labeled points in our maize dataset that describe the position of the leaf or root veins are not specific. They are considered correct as long as they do not deviate from the vein where the leaf or root veins are located, which is different from the keypoints for body used in the human keypoint detection task such as OpenPose (26), where each of the 25 keypoints has precise location constraints. Based on the inference that these “keypoints” share similar characteristics in our dataset, we have modified the traditional method of predicting keypoints. Instead of inferring a separate heatmap for each individual keypoint, we predict all keypoints within an object using a single heatmap. By doing so, we can obtain the positions of the peak values in the heatmap, which serve as the “keypoints” we need. This process is followed by applying NMS, as introduced in Section Soft-NMS, to determine the final locations of the keypoints.

#### Feature pyramid network

The Feature Pyramid Network(FPN) (27) was proposed to address the challenge of detect objects at different scales in an image, which is particularly relevant in our maize dataset where leaves exhibit significant variation in size and shape. FPN, as illustrated in Figure 4, consists of a bottom-up pathway and a top-down pathway. The bottom-up pathway is a standard CNNs that extracts features from the input feature maps. On the other hand, the top-down pathway is responsible for creating a feature pyramid, which is a set of feature maps at different scales based on the input. The top-down pathway takes the highest resolution feature map from the bottom-up pathway and upsamples it to the next scale. Subsequently, the upsampled feature map is merged with the feature map from the bottom-up pathway for that particular scale. This iterative process is repeated for each scale in the pyramid, resulting in a set of feature maps that are leveraged for object detection. FPN has been integrated with Mask R-CNN to further enhance object detection and segmentation performance. In Section Object detection results of maize based on three different original models, we will also present a comparison of our method to other approaches that do not employ FPN, where we demonstrate the superior performance of our proposed technique.

**Fig. 4.**
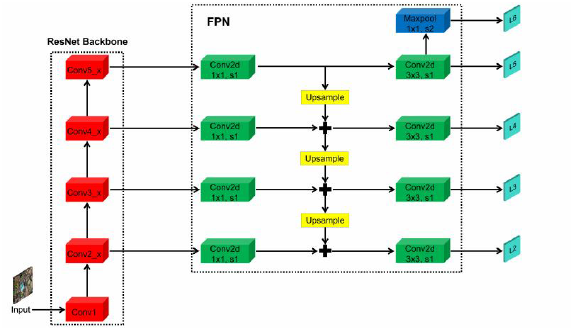
Feature pyramid network structure.

## Results

### Performance evaluation metrics

#### mean Average Precision (mAP)

The mean Average Precision (mAP) is a widely used evaluation metric for object detection algorithms. It servers as a measure of the accuracy and completeness of the detection results. mAP is calculated by averaging the precision scores at various recall levels. Precision represents the ratio of true positive detections to to the total number of detections, while recall represents the ratio of true positive detections to the total number of ground truth objects,. The formulas for precision and recall are as follows:

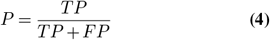

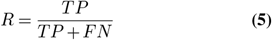

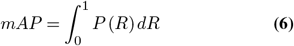

In the first stage, Point-Line Net generates a list of bounding boxes along with their corresponding confidence scores. These confidence scores indicate the likelihood that the object inside the bounding box belongs to a particular class. Subsequently, the detection results are compared to the ground truth objects, which consists of the actual objects in the dataset, including their respective bounding boxes and class labels. For each class, precision and recall values are computed at various confidence thresholds. These values are then used to plot a precision-recall curve. The area under the curve (AUC) is calculated as a measure of performance. The mean Average Precision (mAP) is obtained by averaging the AUC values across all classes. In our study, the mAP metric was evaluated based on the IOU threshold of 0.5, which is a commonly employed threshold for object detection task. A higher mAP value indicates superior detection performance. The mAP metric is widely utilized in object detection benchmarks, including COCO (28), Pascal VOC (29), and ImageNet (30). We also employ mAP as a measure to assess the effectiveness of the object detection stage.

#### Line-Distance

In keypoints detection tasks, standard validation metrics typically rely on the Object keypoint Similarity (OKS)((31), (32)), which is represented as:

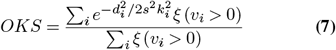

Where *d*_*i*_ is the Euclidean distance between the detected key-point and the corresponding ground truth, *v*_*i*_ is the visibility of the ground truth, *s* is the square root of the area size that is occupied by this object. However, such an evaluation metric, which is applicable to most keypoint detection tasks, is not suitable for our specific task. While our proposed method draws inspiration from keypoint detection, our goal is distinct. We aim to predict whether the keypoints are located on the the “key-path” (leaf or root veins) rather than requiring precise localization of each keypoint. Therefore, even if the predicted keypoints deviate slightly from the exact location, as long as they remain within the correct path, we consider this to be a correct prediction. Based on the aforementioned analysis, we have developed a novel evaluation metric called mean-Line-Distance (mLD) to access the accuracy of key-point detection for our specific requirements. The calculation process of mLD is as follows:

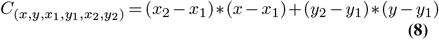

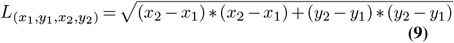

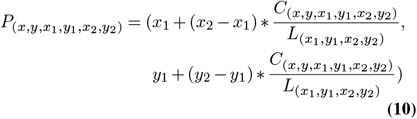

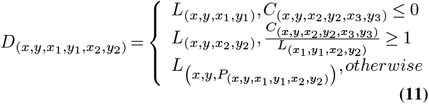

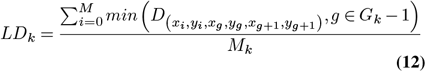

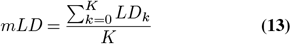

Where *K* denotes the number of detected targets, while *M* represents the number of keypoints that constitute each target after the post-processing operation, *G* refers to the number of keypoints of the ground truth that corresponded to each target. Consequently, *LD*_*k*_ signifies the distance between the *k*_*th*_ target and its corresponding real target, which is obtained by calculating the average distance between the *M* predicted keypoint and the real target. Given that each ground truth is labeled by multiple keypoints, we define the distance from the predicted point to the target as the shortest distance from the predicted point to the line segment formed by these multiple keypoints.

## Experiments and results

### Experimental parameter settings

The training and validation experiments took place at the Agricultural Genomics Institute of the Chinese Academy of Agricultural Sciences, utilizing a computing server equipped with an NVIDIA Tesla V100S PCle GPU with a 32GB memory capacity.. The operating system used was Linux 3.10.0. For deep learning training and validation, Pytorch1.10.1 and Cuda11.1 were utilized, which were specifically tailored to the server environment. As explained in Section Dataset processing, the input images were resized from 6016^*^ 4016(or 4016^*^ 6016) to 1504^*^ 1004(or 1004^*^ 1504). Prior to converting the RGB images to tensor format, random color adjustment and random horizontal flipping were applied to enhance data diversity and improve model robustness. These pre-processing operations, as demonstrated in previous extensive experiments((33), (24), (34)), have proven to be effective in mitigating over-fitting. During the training process, we utilized the Stochastic Gradient Descent (SGD) optimizer with momentum. The model was trained for 200 epochs, commencing with an initial learning rate of 1e-3 and a weight decay of 1e-4. To reduce the learning rate, we employed the StepLR strategy, which involved decreasing the learning rate by a factor of 0.66 every 10 fixed steps.

## Results

### Object detection results of maize based on three different original models

In our experiments, we evaluated the object detection accuracy for the maize dataset using three popular target detection models: Faster R-CNN, RetinaNet and YOLOv3. All three models have the capability to incorporate additional branches for tasks such as keypoint detection or instance segmentation. We did not attempt to modify the default backbone architecture of YOLOv3. While it is technically feasible to replace the backbone with other models like Resnet50 et al, the original Darknet architecture is well-suited for YOLOv3 and generally recommended for optimal performance. Based on intuition, we selected the model that achieved the highest accuracy for further study. The object detection results are presented in Figure 5 and Table 1.

**Table 1.**
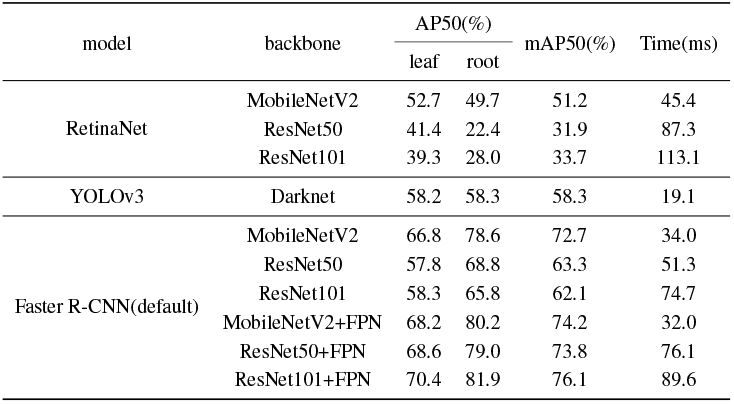
Object detection results for models with different backbones.

**Fig. 5.**
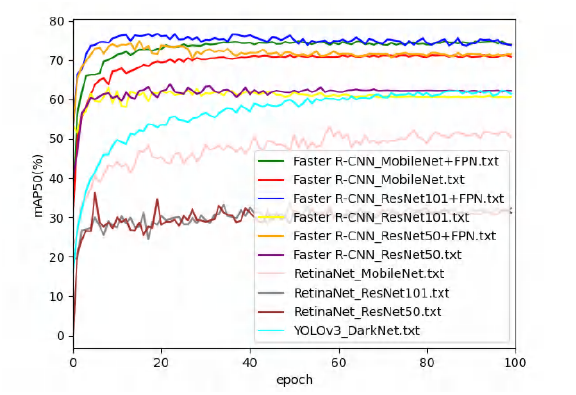
mAP50(%) achieved using different models.

It can be observed that Faster R-CNN (default) with ResNet101 + FPN achieved superior performance compared to other models, with a highest mAP50 score of 76.1% The detection speed, which refers to the time required to detect each image, was measured at 89.6ms. Although this speed is longer than that of other backbones, it remains within a reasonable range that meets we expectations. It is worthy noting that no optimization operations were performed for the results shown in Table 1, and all hyper-parameters were employed as described in the original paper. Therefore, we can consider the selection of the model with the best object detection performance for the subsequent study to be reliable.

## Results

### Object detection results of maize based on improved Faster R-CNN

After selecting Faster R-CNN with ResNet101 + FPN as our object detector, we proceeded to improve the accuracy of object detection. As explained in Section Maize detection, the accuracy of downstream branches, such as key-point detection, is greatly influenced by the performance of the detector in top-down object detection models including Faster R-CNN,. To begin with, we fine-tuned certain hyper-parameters and conducted corresponding experiments. The results of these experiments are summarized in Table 2.

**Table 2.** Object detection results for Faster R-CNN with fine-tuned hyper-parameter.

| Parameter | value | AP50(%) |  | mAP50(%) |
| --- | --- | --- | --- | --- |
|  |  | leaf | root |  |
| RPN_PRE_NMS+RPN_POST_NMS | 2000+1000 | 70.4 | 81.9 | 76.0 |
|  | 3000+1500 | 71.0 | 82.3 | 76.7 |
|  | 4000+2000 | 73.1 | 82.8 | 78.0 |
|  | 5000+2500 | 71.7 | 83.1 | 77.4 |
|  | 8000+4000 | 72.8 | 80.2 | 76.5 |

The results demonstrate that the model performed optimally when the number of retained proposals in RPN before and after NMS processing was set to 4000 and 2000, respectively. Additionally, we made further adjustments as outlined in Section Adjustment of hyper-parameters. Apart from the aforementioned hyper-parameters, we discovered that other factors, such as aspect_ratios of the anchor generator and the IoU thresholds used in NMS processing in the RPN, had minimal impact on the detection accuracy during our experiments. Hence, we followed the settings specified in the original paper for these parameters. In order to address the issue of target occlusion present in the dataset, we incorporated Soft-NMS and D_IoU techniques. The experimental results are presented in Table 3:

**Table 3.** Object detection results for Faster R-CNN with Soft-NMS and D_IoU.

| NMS | Soft-NMS | D_IoU | AP50(%) |  | mAP50(%) |
| --- | --- | --- | --- | --- | --- |
|  |  |  | leaf | root |  |
| ✓ |  |  | 73.1 | 83.1 | 78.0 |
|  | ✓ |  | 75.1 | 85.4 | 80.2 |
|  |  | ✓ | 74.9 | 84.4 | 79.7 |
|  | ✓ | ✓ | 75.5 | 86.0 | 80.8 |

The results obtained from the above-mentioned experiments are in line with our initial hypothesis. Considering the significant occurrence of maize leaf occlusion in our dataset, particularly during the later stages of growth as depicted in Figure 1, many targets are closely positioned or occluded. Consequently, when performing NMS processing, the model may mistakenly filter out genuine targets. However, with the incorporation of Soft-NMS and D_IoU techniques, these ground-truth targets are retained to a greater extent, leading to an improvement of 2.8% in the mAP50 of the model.

### Vein recognition results with Point-Line Ne

Inspired by human keypoint detection, we initially attempted to utilize traditional keypoint detection methods to identify leaf and root veins of maize. As the experiment progressed, we made innovative improvements described in Section Modification of keypoint detection branch. The results of the comparative experiments for the two “keypoint” detection methods are presented in Figure 6 and Table 4:

**Table 4.** Leaf vein and rootstocks vein detection results using different method.

| Method | AP50(%) |  | mAP50(%) | Line-Distance |
| --- | --- | --- | --- | --- |
|  | leaf | root |  |  |
| Traditional method | 75.5 | 86.0 | 80.8 | 34.1 |
| Innovative method | 76.5 | 86.6 | 81.5 | 33.5 |

**Fig. 6.**
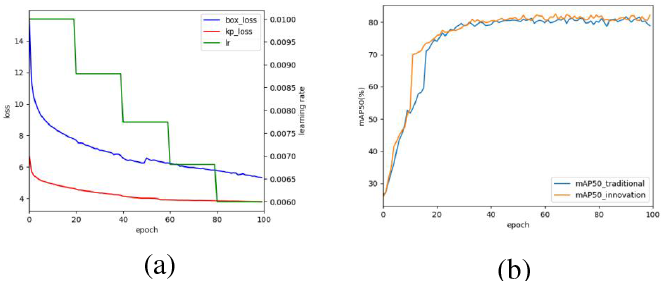
Training curves: (a) loss curve and learning rate curve changes of traditional and innovative methods, where box_loss consists of RPN Loss and Fast R-CNN Loss (7), and kp_loss refers to the cross entropy loss of the keypoint detection branch; (b) mAP50 curve changes of traditional and innovative methods.

The experimental results indicate that our innovative method achieves a smaller distance (measured using custom Line-Distance) between the predicted keypoints of the leaf or root and the ground-truth veins. Specifically, the distance is reduced to 33.5. This observation further demonstrates that the improvements we have implemented provide better perfor-mance in accurately describing the position of the leaf and root veins according to our requirements. It is worth noting that the magnitude of this metric is related to the scale size of the input image.

### Training performance record with Point-Line Net

Through-out the training and validation process, we carefully adjusted the parameters and monitored the performance of model to identify signs of overfitting or underfitting. We recorded the changes in learning rate(lr), various losses and evaluation metrics, as depicted in Figure 5. The graph illustrates that as the model was trained up to the 100_*th*_ epochs, the mAP tended to stabilize and would not increase significantly. Although the total loss continued to decrease gradually, there was evidence of overfitting. Consequently, we saved the best weights obtained after the 100_*th*_ epoch as the final result of model training for subsequent prediction tasks. To showcase the performance of our model, we selected several images from the validation set that presented varying levels of complexity across different growth stages. We then compared the predicted results with the ground-truth annotations. The visualization results are presented in Figure 7. Notably, as the complexity of the background environment increased and occlusion between targets became more prevalent, the tasks of object detection and keypoint detection became progressively challenging. Nonetheless, our model exhibited commendable performance in such scenarios.

**Fig. 7.**
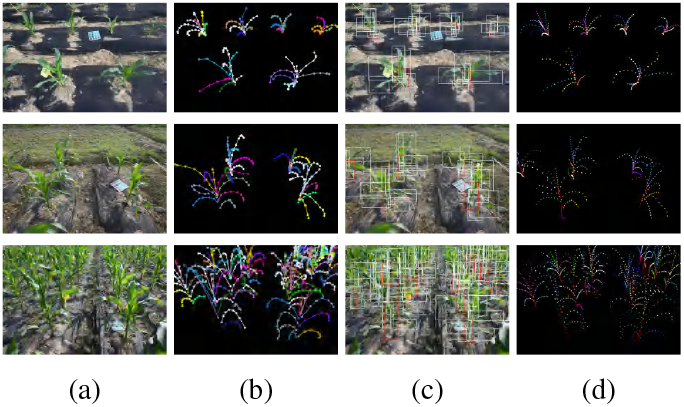
Prediction results of Point-Line Net in various scenarios: (a) Original RGB images; (b) Ground-truth transformed by labelme tool; (c) Predicted bounding boxes; (d) Predicted keypoints.

## Conclusions

Deep learning methods have gained significant popularity in the field of plant phenotyping due to their exceptional capability in feature extraction and classification. Plant phenotyping aims to quantify various traits of plants, including the size, shape, color, texture, and growth rate of leaves, stems, and flowers. These traits serve as crucial indicators of plant growth, development, and stress responses. They can further facilitate the selection and breeding of superior crops, enhance crop management practices, and predict crop yields. Deep learning methods excel in automatically learning complex and high-dimensional features from plant images, eliminating the need for time-consuming and subjective manual feature engineering. Among these methods, CNNs are particularly renowned for their ability to learn spatial features across multiple scales and orientations from raw images. From the inception of our maize dataset, our team has been dedicated to utilizing deep learning methods for field crop phenotype detection. To the best of our knowledge, there is currently no field crop phenotype dataset that employs our innovative dotted-line annotation approach to represent objects, such as leaves or stems, nor is there a dataset as large as ours. Nevertheless, this innovative annotation method, coupled with the complex field environment, and the random growth pattern of maize, presents significant challenges in achieving accurate detection. Prior to implementing the method proposed in this paper, we attempted various approaches, to detect maize leaves using our maize datatset, including instance segmentation. However, none of these methods yielded satisfactory results until we connected this task with human pose estimation, also known as multi-person keypoint detection. As the original annotation information lacked ground-truth bounding boxes, we naturally experimented with the bottom-up keypoint detection method described in Section Point-Line Net Based Mask R-CNN. Unfortunately, the performance was still inadequate. We assume that handing the clustering problem for multiple targets in complex scenarios, particularly with significant overlap, proved to be challenging. Based on our previous explorations, we decided to adopt a top-down key-point detection scheme that incorporated the use of bounding boxes to meet more accurate detection requirements. Subsequently, we fine-tuned the parameters, selected the optimal backbone, and introduced numerous innovative improvements. Each optimization was accompanied by rigorous experimental testing. Ultimately, we achieved an mAP50 of 81.5% and a custom distance evaluation index (LD) of 33.5, which holds significant implications for field maize phenotype detection. Upon reviewing the work presented in this paper, we acknowledge that there is still consider room for improvement. Firstly, our proposed method remains limited by bounding boxes, contrary to the annotation approach employed by our team to create the maize dataset. Consequently, the accuracy of the final keypoint detection is constrained by the accuracy of the initial stage object detection, as mentioned Section Point-Line Net Based Mask R-CNN. In future research, We aspire to achieve the task of detecting leaf and root veins without the aid of bounding boxes. Secondly, since there is currently no standard evaluation metric to assess the accuracy of predicted keypoints, we devised the customized index LD. However, there is room for refinement in order to enhance the comprehensiveness and precision of the evaluation. Additionally, it is important to highlight that our study exclusively employed field images captured by cameras and a fraction of the annotation information provided in our maize dataset. Moving forward, we aim to conduct more in-depth exploration by incorporating other valuable annotation information, such as leaf attribution information, multi-angle location information, and additional images captured by UAVs.

## ACKNOWLEDGEMENTS

**Jue Ruan, Dengao Li**: Methodology, Conceptualization, Resources, Funding acquisition, Supervision. **Bingwen Liu, Dengfeng Hou, Jianye Chang**: Investigation, Data curation, Formal analysis, Writing – original draft. **Jianye Chang**: Resources, Validation, Writing – review & editing.

